# Universal Transcriptional Responses to Different Modalities of Traumatic Stress as a Measure of Stress Exposure

**DOI:** 10.1101/185926

**Authors:** Moriah L. Jacobson, Lydia A. Kim, Barbara Rosati, David McKinnon

**Affiliations:** Department of Psychology, Stony Brook University, Stony Brook, NY 11794, U.S.A; Department of Physiology and Biophysics, Stony Brook University, Stony Brook, NY 11794, U.S.A; Institute of Molecular Cardiology, Stony Brook University, Stony Brook, NY 11794, U.S.A; Department of Veterans Affairs Medical Center, Northport, NY 11768, U.S.A.

**Keywords:** gene regulation, transcriptome, stress, adrenal gland, HPA axis

## Abstract

Molecular diagnostic tools that can robustly and quantitatively measure the response to traumatic stress would be of considerable value in assessing the individual risk of developing post-traumatic stress disorder or stress-induced depression following stress exposure. The gene regulatory network can integrate and encode a large number of different signals, including those elicited by exposure to stress. We find that many genes respond to at least one modality of stress but only a subset of stress-sensitive genes track stress exposure across multiple stress modalities and are thus universal markers of stress exposure. A sensitive and robust measure of stress exposure can be constructed using a small number of genes selected from this modality-independent set of stress-sensitive genes. This stress-sensitive gene expression (SSGE) index can detect chronic traumatic stress exposure in a wide range of different stress models in a manner that is relatively independent of the modality of stress exposure and that parallels the intensity of stress exposure in a dose-dependent manner.

## Introduction

Exposure to traumatic stress can lead to the development of post-traumatic stress disorder (PTSD) and/or stress-induced depression, both of which can cause severe personal distress as well as interfere with occupational and social function, resulting in significant individual and social costs (1). There is considerable interest in developing molecular diagnostic tools for the assessment of traumatic stress exposure (2). Ideal markers for traumatic stress exposure would be graded, or dose-dependent, producing an increasing signal with increasing intensity of stress exposure thereby allowing quantitative assessment of the degree of stress exposure. A second requirement is that the diagnostic test be robust, meaning that the test would be similarly responsive to the broad range of different traumatic stressors that an individual might encounter.

The effects of traumatic stress exposure are encoded in both the gene regulatory and neural networks of individuals subjected to stress (3-5). One approach to developing molecular diagnostic tools is to use changes in gene regulatory function as markers of stress exposure (6). The gene regulatory network integrates a large number of different signals in order to yield a particular pattern of gene expression (7). In this study, we sought to find those nodes in the regulatory network that are the most sensitive and robust indicators of stress exposure.

There is no single, universally agreed upon pre-clinical model of PTSD or stress-induced depression. This is because a wide variety of stressors and temporal patterns of stress exposure can produce long-lasting changes in psychological and physiological function (8, 9). In general, studies on the development of molecular markers for stress exposure have used only a single stressor paradigm to elicit a stress response. This approach carries the risk that the biomarkers identified in these studies will only be representative of that particular stress paradigm and will not adequately characterize responses to the broad range of stress paradigms that are used experimentally or, more importantly, to the diverse set of traumatic stressors that can occur in life. A primary purpose of this study was to determine the feasibility of identifying biomarkers that respond to most, or all, stress modalities.

In this report, RNA sequencing was used to examine the response of the transcriptome to a variety of different chronic stress protocols. Changes in gene regulatory function were examined in the adrenal gland, which plays a pivotal role in mediating the stress response, as both the terminal organ in the HPA axis as well as a major component of the peripheral sympathetic nervous system (10, 11). We reasoned that, if a universal response to chronic stress could not be observed in this tissue, it would be unlikely to be present in other tissues, which have more diverse physiological roles, independent of the response to stress. Large differences in the response to different stress protocols were observed, as measured by changes in gene expression, but a set of stress modality independent responses, which may reflect a universal response to chronic stress in this tissue, were also observed. Members of this set of universal response genes were used to construct a stress sensitive gene expression index that was capable of reliably detecting and quantitating differences in traumatic stress exposure across six different chronic stress models.

## Methods and Materials

### Animals

Male Sprague-Dawley rats were used in all experiments. All procedures were approved by the Institutional Animal Care and Use Committee (IACUC) at Stony Brook University.

### Chronic Stress Models

Seven different animal treatment protocols were used. Protocols were typically 3 weeks in duration and started at postnatal day 28. More detailed descriptions of these protocols are given in the Supplemental Information.

#### Control (C)

Animals lived socially in groups of three per cage, with no additional stressors other than daily weighing and routine husbandry.

#### Social Isolation (SI)

Animals were singly housed, with no additional stressors other than daily weighing and routine husbandry.

#### Social Defeat (SD)

Animals were exposed to daily sessions of defeat in the home cage of male Long Evans rats. Several variations of defeat were used including direct physical defeat, psychological threat of defeat, and witnessing defeat in a conspecific. For socially housed defeat, the rats were housed socially in groups of three per cage, before and after the defeat sessions.

#### Isolation Defeat (ID)

This protocol was identical to the Social Defeat protocol except that the rats were singly housed, before and after the defeat sessions, for the duration of the stress protocol.

#### Grid housing (GH)

Animals lived singly on metal grid of shock apparatus (Coulbourn Instruments), with no additional stressors other than daily weighing and routine husbandry.

#### Chronic shock (CS)

Animals lived singly on metal grid of shock apparatus and were administered an electric foot shock of randomly varying duration and intensity, at random time intervals.

#### Chronic variable stress (CVS)

Animals were exposed to a series of diverse physical, psychological and psychosocial stressors, including predator threat using either live animals or predator scent, water submersion, cold and warm room exposure, cage tilt and rotation, restraint, bedding disruptions, circadian rhythm disruption, shock, food and water deprivation, forced swim, isolation, and social instability.

## RNA Preparation, Sequencing, Data Analysis and Real-time PCR

Total RNA was extracted from tissue samples using the RNeasy Miniprep Kit (Qiagen). cDNA libraries were prepared using the NEBNext Ultra II Directional RNA Library Prep Kit (New England BioLabs). Alignment of RNA sequencing data was performed using the Rsubread package (12). Count matrices were created using the featureCounts function in Rsubread (13). Raw and processed RNA sequencing data have been deposited in NCBI’s Gene Expression Omnibus database (GSE100454). Real-time PCR analysis was performed using standard methods (14). See Supplemental Information for details.

Raw and processed RNA sequencing data have been deposited in NCBI’s Gene Expression Omnibus database (GSE100454).

## Behavioral Tests

The acoustic startle response test (ASR) and elevated plus maze test (EPM) were performed using essentially standard procedures (15-18). See Supplemental Information for details.

## Statistics

For comparison of multiple means, ANOVA with post-hoc t-tests using Benjamini-Hochberg correction (19) were performed using R (20). Principal component analysis was also performed using R. Effect sizes are reported as Cohen’s d, calculated using 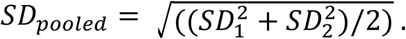

## Results

### Diverse and Common Changes in Response to Intense Chronic Stress

A total of six different chronic stress paradigms were developed: social isolation, social defeat, social defeat with social isolation, grid housing, chronic shock and chronic variable stress (see Methods and Materials for details). We initially focused on the two most intense protocols, chronic shock (CS) and chronic variable stress (CVS), which use very different methods to induce stress.

In the adrenal gland, a relatively large number of genes were found to be differentially expressed following exposure to each of these protocols (Figure 1A). There were more changes in response to the chronic shock protocol than to the chronic variable stress protocol, with 196 genes differentially expressed following the chronic shock protocol versus 124 genes following the chronic variable stres protocol (FDR = 0.05). Notably, a majority of the genes that changed in response to chronic variable stress did not change in the chronic shock animals and vice versa (Figure 1B).

**Figure 1.**
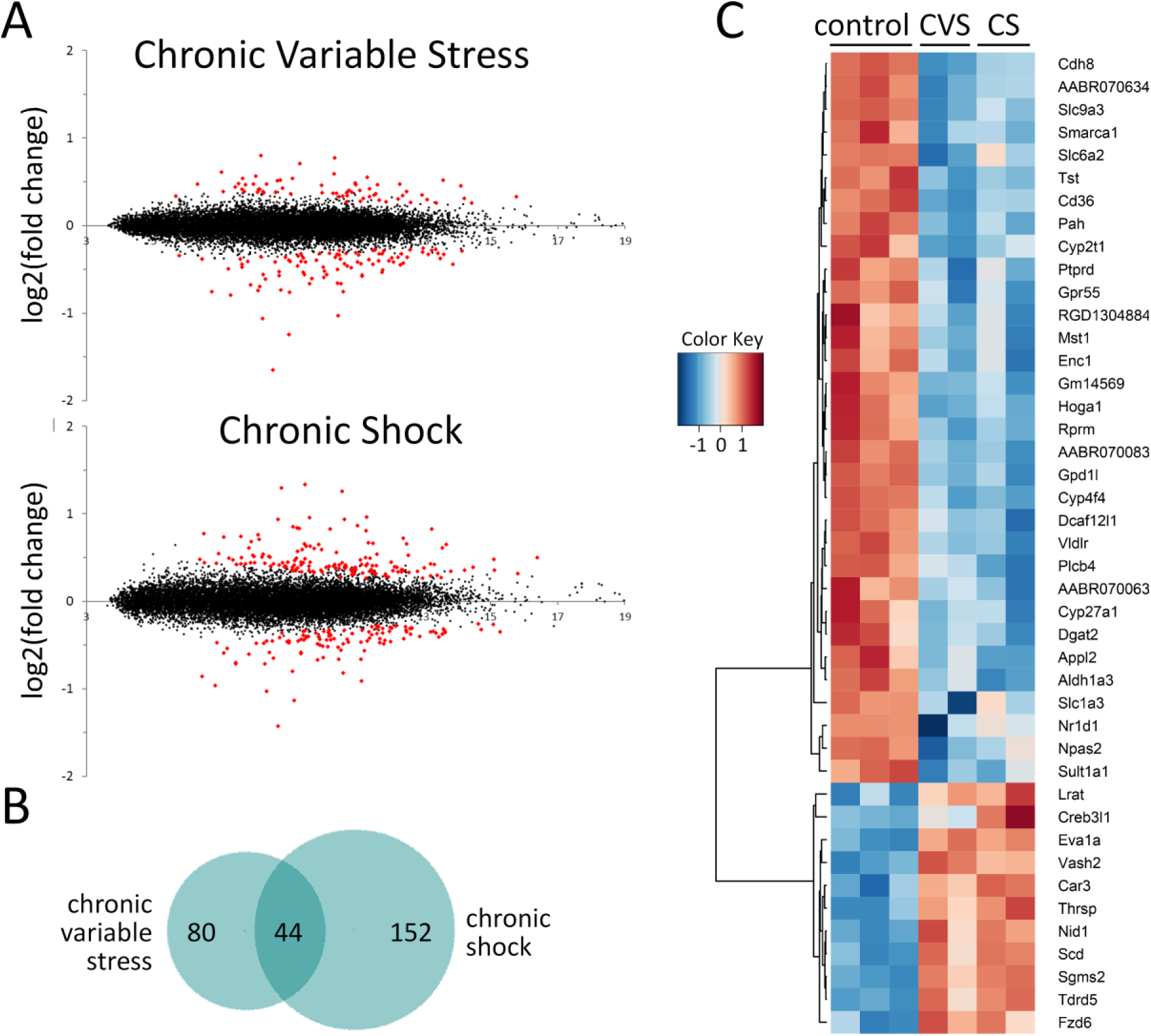
A plots of RNA sequencing data from adrenal gland for animals exposed to the chronic variable stress (CVS) or chronic shock (CS) protocols. The x-axis corresponds to log2(average expression) and the y-axis to log2(Stress/Control). Differentially expressed genes (marked in red) were selected using FDR = 0.05. RNA samples were pooled (n=8-9) before sequence analysis for each of the four independent replicates of the experimental groups (2 CVS and 2 CS) and three independent control groups. Fold-change values reflect dispersion moderation in DESeq2. See Tables S2 and S3 for complete list of differentially expressed genes. B. Venn diagram of the differentially expressed genes for the two different stress groups. The total number of differentially expressed genes was 124 for the CVS protocol, 196 for CS protocol and 44 genes were differentially expressed in both protocols. C. Heat map for those genes changed in both CVS and CS stress protocols. Expression values were log2 transformed and scaled to each row.

A subset of 44 genes was found to be differentially expressed in both protocols (Table S1). With a few exceptions, this common set of differentially expressed genes changed similarly in response to both stress protocols (Figure 1C). A preponderance of these genes were down-regulated. Gene ontology analysis did not point to any particular process or processes in which this common set of genes were broadly involved.

### Stress-Sensitive Gene Expression Index

The existence of a common set of differentially expressed genes, which respond to both of these very different stress protocols, suggested that these genes might be universal markers of traumatic stress exposure in the adrenal gland. We hypothesized that a subset of these common response genes could be used to construct a stress-sensitive gene expression (SSGE) index that could be used to measure stress exposure in a consistent and quantitative way across a broad array of chronic stress modalities. It was anticipated that this index might function in an analogous way to a stock market index, which can capture the mood of the market by sampling only a small subset of the most characteristic stocks.

Genes were selected from the common set of 44 genes for potential inclusion in the stress-sensitive gene expression index based on several criteria. Genes with the largest effect size and for which the fold-changes were similar in both the CS and CVS protocols were selected. In addition, genes with consistent expression levels within the control group, as measured using real-time PCR, were favored. Based on these criteria, six genes were selected: *Pah*, *Slc9a3*, *Thrsp*, *Scd*, *Cdh8* and *Cd36*. The index was deliberately restricted to a relatively small number of genes in order to simplify measurement across large numbers of individual animals.

Principal component analysis (PCA) of the expression values derived from real-time PCR analysis of gene expression for the six genes selected for the index demonstrated that there was no overlap between individuals in the stress (CVS or CS) and control groups along the axis of the first principal component (Figure 2A). This was important because it implies that these expression values could be used to reliably distinguish individuals exposed to chronic stress from controls, essentially functioning as a diagnostic test.

**Figure 2.**
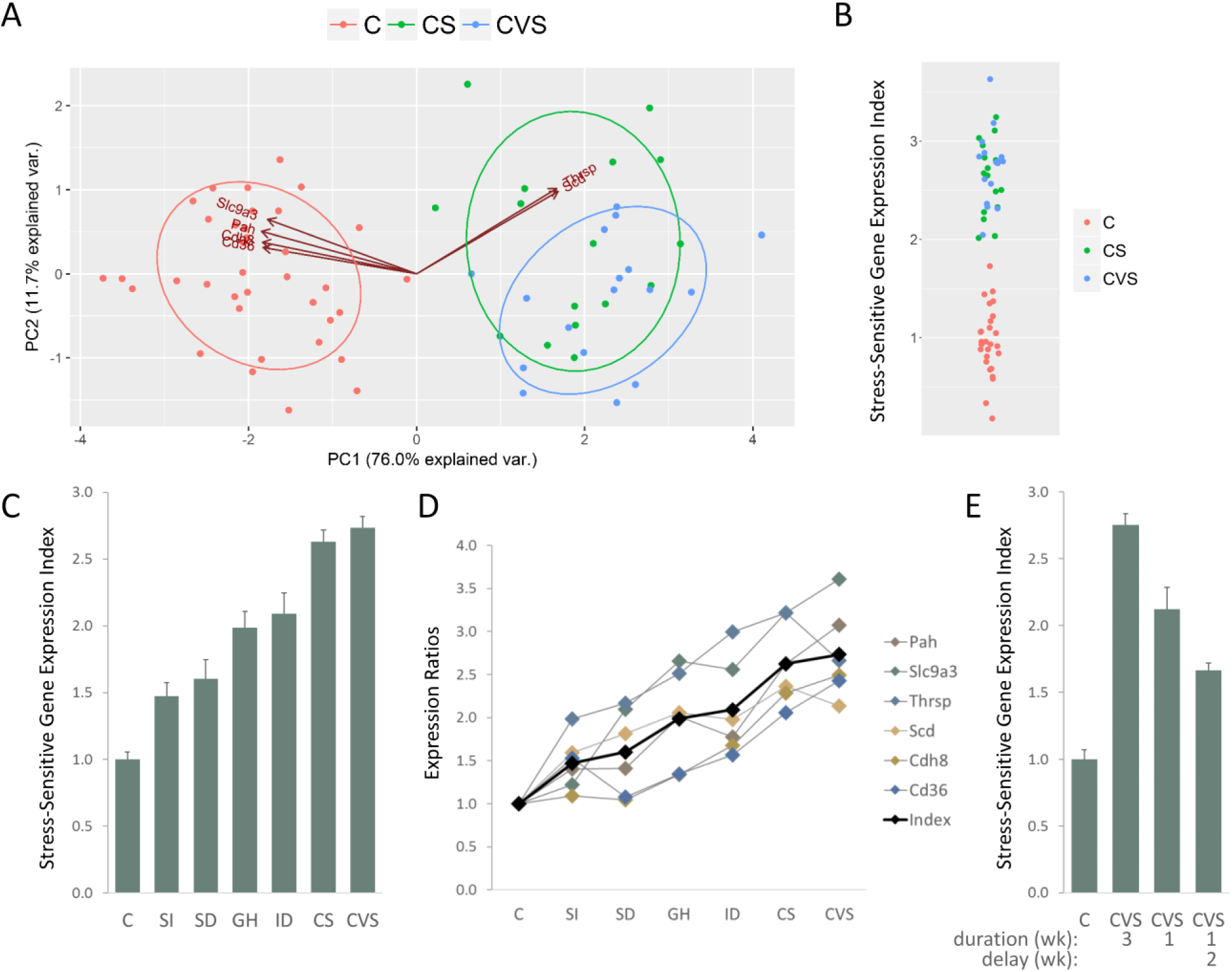
A. Principal component analysis using real-time PCR expression values for six genes (*Pah*, *Slc9a3*, *Thrsp*, *Scd*, *Cdh8*and *Cd36*) comparing control (n=36) animals and animals exposed to the two most intense stress protocols (n=33), either chronic variable stress (CVS) or chronic shock stress (CS). B. Strip plot comparing the stress-sensitive gene expression index for control animals and animals exposed to either chronic variable stress (CVS) or chronic shock stress (CS). C. Average stress-sensitive gene expression (SSGE) index for control (C) and six different chronic stress groups: social isolation (SI), social defeat (SD), grid housing (GH), isolation defeat (ID), chronic variable stress (CVS) and chronic shock (CS). There was a statistically significant difference between the groups as determined by one way ANOVA (F(6,105) = 55.97, p < 2×10^−16^). All stress groups were different to the control group: SI (p = 1×10^−4^, d = 1.2), SD (p = 1×10^−4^, d = 1.5), GH (p = 1×10^−8^, d = 2.8), ID (p =6×10^−10^, d = 2.8), CS (p < 2×10^−16^, d = 4.5) and CVS (p < 2×10^−16^, d = 4.9). D. Mean expression ratios for each of the six selected genes compared to the index (black) across the seven different experimental groups. E. Comparison of SSGE index for control (C), the standard 3 week duration CVS protocol (CVS), a protocol of 1 week CVS and a protocol of 1 week CVS followed by 2 weeks of social housing. There was a statistically significant difference between the groups as determined by one-way ANOVA (F(3,48) = 76.5, p < 2×10^−16^). All stress groups were different to the control group: CVS (p < 2×10^−16^, d = 5.3), CVS 1 week (p = 2×10^−9^, d = 2.9) and CVS 1 week with 2 week delay (p = 6×10^−3^, d = 2.7). Error bars are s.e.m.

A Stress-Sensitive Gene Expression (SSGE) index was then calculated as follows. For each individual, *j*, an expression ratio, 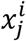was determined for each gene, *i*,

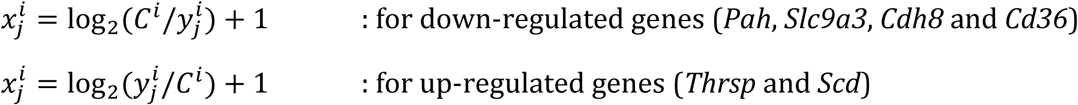

where,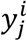was the mRNA expression value for gene, *i* in individual, *j* and *C^i^* was the mean expression level for gene, *i* in the control animals. Expression ratios were converted to logarithm base 2 to maintain within group variance relatively constant across the different experimental groups. The expression ratios, 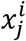, were normalized so that the mean, 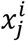, for the control samples, for each gene, *i*, was equal to unity. Expression ratios 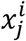for each of the genes were then averaged to give the SSGE index, I_*j*_, for each individual, *j*. The index has a mean value of unity for the control samples.

The SSGE index performed as well as principal component analysis in distinguishing all the individual animals in the CVS and CS groups from the control animals (Figure 2B). The advantage of the index over PCA was that the index scale is only dependent on the average expression values in the control samples. These average values were stable across different experimental replicates, facilitating the comparison of stress exposure across different protocols.

### Stress-Sensitive Gene Expression Index Applied to Multiple Stress Protocols

To determine whether the index could perform in a predictable manner across different modalities of stress exposure, average index values were determined for the control group and the six different stress protocol groups (Figure 2C). There was a statistically significant difference between the groups as determined by one-way ANOVA (F(6,105) = 55.97, p < 2×10^-16^) and all stress groups were different to the control group.

The average SSGE index values increased in a manner consistent with the relative intensity of the different stress protocols. Social isolation was the least stressful of these protocols but exposure to this protocol could still be readily detected, with the SSGE index being higher in the stress group than controls, who were socially housed (p = 1×10^-4^, d = 1.2). It is reasonable to assume that grid housing (GH) would be more stressful than social isolation since it combines social isolation with difficult housing conditions. Indeed, the SSGE index was found to be higher for grid housing in comparison with the social isolation group (p = 3×10^−3^, d = 1.3). Similarly, it is reasonable to assume that chronic shock would be more stressful than grid housing, since this adds electric shocks to the social isolation and difficult housing conditions. As expected the index was higher for this group relative to grid housing (p = 5×10^−4^, d = 1.8). The relatively high score for grid housing was consistent with our observations of the rat’s general condition and the lengths to which the animals would go to get off the grid, even in the absence of shock.

Based on body and organ weight changes (see below), the chronic variable stress (CVS) and chronic shock (CS) stress protocols were more stressful than the other protocols and were similar in intensity to each other. This assessment was also reflected in the stress index results (Figure 2C). Rats are less likely to display symptoms of anxiety following social defeat when they are housed socially rather than in isolation (21), suggesting that the isolation defeat (ID) is more stressful than social defeat (SD). The difference in the respective index values for the two protocols (p = 0.01, d = 1.1) was consistent with this expectation.

Although the individual expression ratios for each of the six genes that contribute to the index responded somewhat differently to the different stress protocols, the trend for each expression ratio across the six different protocols was in broad agreement with the overall trend of the SSGE index (Figure 2D). For the three most intense protocols (ID, CS and CVS) all six individual genes were differentially expressed relative to the controls (p < 0.005). These genes can reasonably be considered universally responsive to these different stress modalities. The number of genes that were differentially expressed relative to controls fell as the protocols became less intense (GH: 5/6, SD: 3/6, SI: 3/6), suggesting that some genes in the index are more responsive to lower levels of stress than others.

The performance of the index was not particularly sensitive to the number or combination of genes included in the index. Reducing the number of index genes to only four, those with the largest effect size (*Pah*, *Slc9a3*, *Thrsp* and *Scd*), did not substantially alter the performance of the index, although it did result in a modest increase in the within-group variance (average S.D. for each group with six index genes was 0.39 versus 0.49 with only four genes).

To examine the effect of different durations of stress exposure, the standard CVS protocol (3-week duration) was compared with a shorter CVS protocol (1-week duration) (Figure 2E). The magnitude of the SSGE index after the 1-week stress exposure was smaller than that seen for the 3-week protocol (p = 1×10^−4^, d = −1.5), but remained well above control levels (p = 2×10^−9^, d = 2.9), suggesting that shorter periods of stress exposure can also be readily detected by the index. The time period over which the SSGE index remains elevated following the cessation of stress exposure was examined by exposing animals to a 1-week duration CVS protocol and then returning them to social housing for two weeks before analysis. In this case the SSGE index remained elevated compared to controls (p = 6×10^−3^, d = 2.7) although the index value was smaller than for the other two CVS groups, where stress exposure occurred closer to the time of measurement (Figure 2E). This result suggests that the index decays relatively slowly, with a time period of weeks.

### Weight Changes in Response to Chronic Stress

Reduction in body weight is a common response to chronic stress exposure (22). Changes in body and organ weight in response to the different stress protocols were consistent with the results from the SSGE index suggesting that the chronic shock (CS) and chronic variable stress (CVS) protocols were more intense than the other stress protocols (Figure 3). Average body weight was reduced relative to controls one day after the end of the three-week stress exposure period for both the CS and CVS groups (p = 1×10^−7^, d = −1.6; and p = 1×10^−12^, d = −3.3; respectively) (Figure 3A).

**Figure 3.**
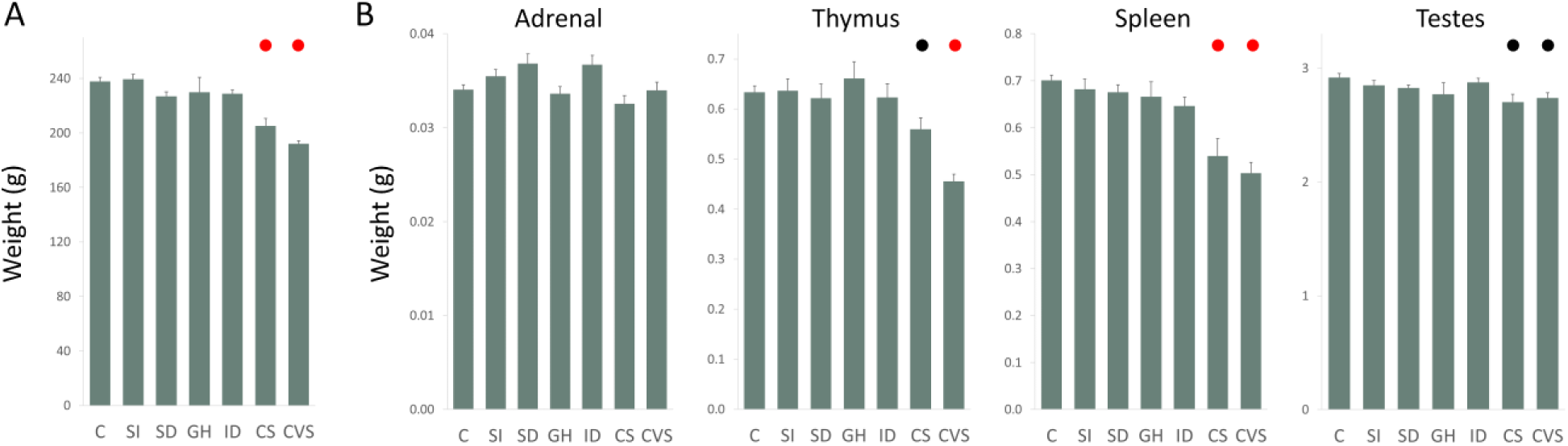
A. Body weight one day after the end of the stress protocols in control and chronically stressed animals. There was a statistically significant difference between the groups as determined by one-way ANOVA (F(6,105) = 18.0, p = 3×10^−14^). Average weights for stress groups that were changed relative to the control group are marked with colored circles, p < 0.05 (black) and p<0.001 (red). B. Organ weights for adrenal gland, thymus, spleen and testes. One-way ANOVA for: adrenal gland (F(6,105) = 2.95, p = 0.01), thymus (F(6,105) = 11.2, p = 1×10^−9^), spleen (F(6,70) = 11.5, p = 6×10^−9^) and testis (F(6,105) = 2.78, p = 0.02). Error bars are s.e.m.

Adrenal gland weight was not changed in any of the stress protocols relative to controls (Figure 3B). Thymus weight was reduced relative to controls for the CVS group (p = 2×10^−9^, d = −2.7) with a smaller effect for the CS group (p = 0.01, d = −0.9). Thymus weight was lower in the CVS group than in the CS group (p = 2×10^−3^, d = −1.3). Spleen weight was reduced for both the CS (p = 4×10^−6^, d = −1.9) and CVS (p = 1×50^−8^, d = −3.2) groups. Testis weight was modestly reduced for both the CS and CVS groups (p = .02, d = −0.9; and p = 0.05, d = −0.9; respectively).

### Behavioral Responses to Chronic Stress

It is has previously been reported that behavioral responses can be quite sensitive to the modality of stress exposure (23) and we observed similar effects in response to our stress models. An example of this is shown for the acoustic startle response (ASR, Figure 4A). In this experiment, ASR was tested before the start of the stress protocol and then tested once a week for the following three weeks of stress exposure. Animals in the chronic variable stress (CVS) and chronic shock (CS) exposure groups responded quite differently in this test (Figure 4A), despite the fact that both stress protocols appear to be similarly intense. Chronic shock exposure produced a persistent decline in the startle response, whereas chronic variable stress had no significant effect on the startle response.

**Figure 4.**
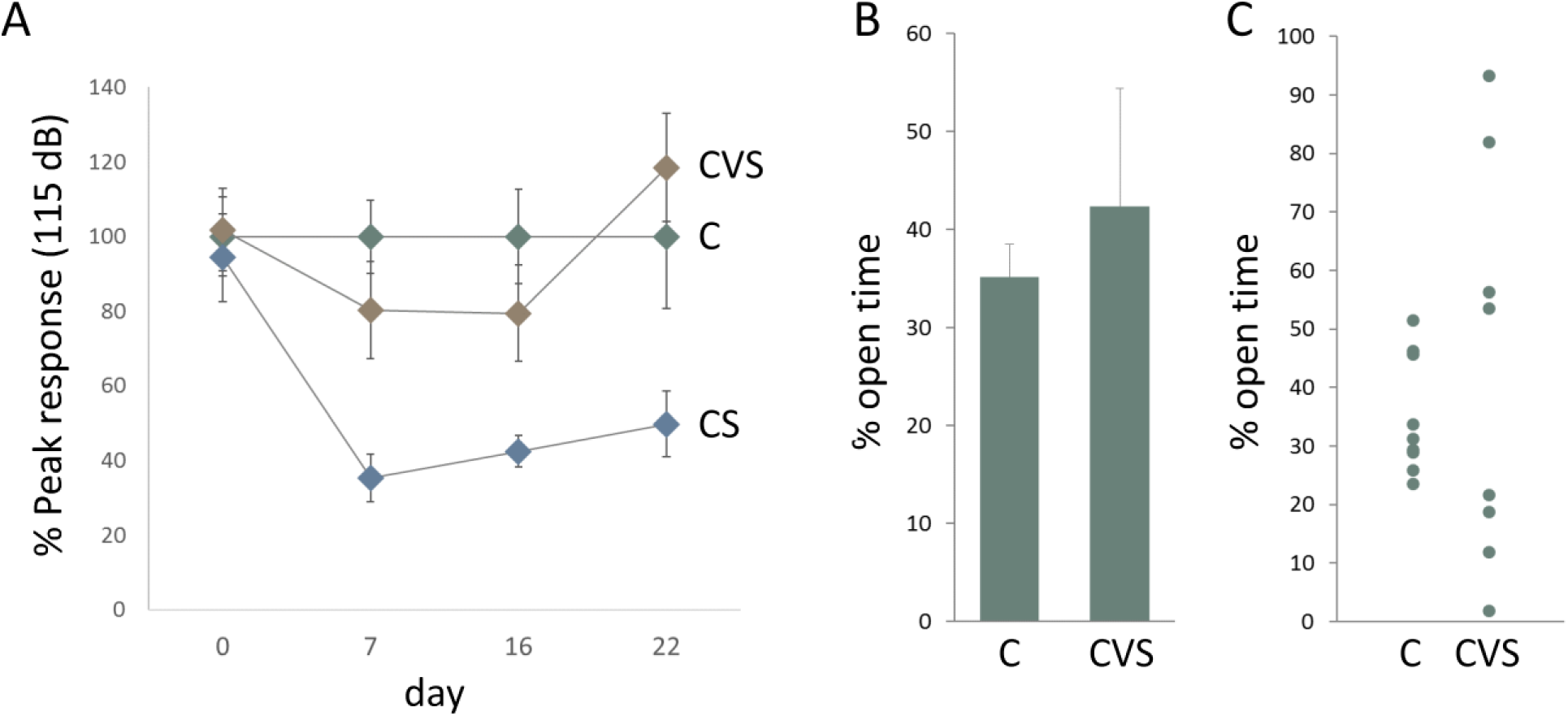
A. Acoustic startle response (ASR). ASR tests were administered on days 0, 7, 16 and 22, relative to the start of the stress protocol on day 1. Results for control (C) (green symbol), chronic variable stress (CVS) (brown symbol) and chronic shock (CS) (blue symbol) are shown. The average startle response of the CS group was lower than the control group on days 7, 16 and 22 (p = 0.0004, d = −2.3; p = 0.002, d = −1.9; and p = 0.01, d = −1.4, respectively) whereas the response of the CVS groups was unchanged on any test day. Data were normalized so that the average peak response of the control group to a 115 dB pulse for each trial was equal to 100. Similar results were also seen using 95 and 105 dB test pulses. Data values are means with s.e.m. error bars (n = 8 or 9). B. Elevated plus maze (EPM). Average time spent in the open arm as a percentage of the total time, for the control and CVS groups. Data values are means with s.e.m. error bars (n = 8 or 9). C. Individual open time values for the same EPM experiment as shown in B.

Modality specific effects were also seen using the elevated plus maze (Figure 4B and C). There was no significant difference between the CVS and the control groups in the mean values for time spent in the open arms of the elevated plus maze (Figure 4B) but there was a large increase in the variance for the CVS group (Figure 4C). The CVS animals tended to ‘freeze’ in one of the compartments. Mean time spent in the ‘frozen’ state over the 5-minute test period was much higher for the CVS animals (92 ± 34 seconds) than for controls (0.8 ± 0.6 seconds) (p = 0.01, d = 1.4). Freezing behavior was also reflected in an average 38% reduction in distance travelled by the CVS group compared to controls (p = 0.009, d = −1.5). In contrast, the chronic shock protocol had no significant effect on the rats’ performance in the elevated plus maze.

## Discussion

In this report, we demonstrate that a subset of stress-sensitive genes responds to traumatic stress exposure in manner that is relatively independent of the modality of stress exposure. A sensitive and robust measure of traumatic stress exposure can be constructed using a small number of biomarkers chosen from this set of genes. This stress-sensitive gene expression index can detect traumatic stress exposure in a wide range of different stress models in a manner that is both relatively independent of the modality of stress exposure and parallels the intensity of stress exposure in a dose-dependent manner.

The data described in this report are consistent with the hypothesis, derived from psychological studies of chronic traumatic stress exposure (24, 25), that the response to traumatic stress exposure is dose-dependent. It is difficult to draw a direct correlation between animal studies and the experiences of humans subjected to extreme and repeated traumatic stress (24-27). Nonetheless, our results are consistent with the hypothesis, developed from these studies, that there is a graded response to stress exposure. In this model, no individual has an absolute resilience to stress, only relative resilience that can be overcome with a sufficiently high ‘dose’ of traumatic stress. Consistent with this idea, every individual animal exposed to the two most intense stress protocols is distinguishable from the control animals, using the stress-sensitive gene expression index (Figure 2B). Similarly, there is no obvious threshold for the effects of stress exposure. The response to even the mildest stressor tested, social isolation, could be detected with this same index.

In addition to the universal stress-response genes, many stress-responsive genes were sensitive to the modality of stress exposure. This suggests that studies using preclinical models should take account of the fact that the stress response, even at the level of transcription, can be quite sensitive to the modality of stress exposure and that validation of any potential stress-sensitive biomarkers requires testing against multiple different models of stress exposure to ensure cross-modality validity.

The index-based measure of stress-sensitive gene expression described in this report is a practical tool for quantitating level of traumatic stress exposure in animal models. It is relatively easy and inexpensive to implement and many steps can be automated so that it can be scaled for large numbers of animals. There are many different ways to measure the effects of chronic stress exposure in laboratory animals (9), including behavioral tests and analysis of endocrine function. Behavioral tests can be very sensitive but behavioral responses can be dependent on the modality of stress exposure (23) (see Figure 4). Endocrine responses to stress can be complex, transitory and habituate in the maintained presence of the stressor (8, 28), and can also depend on stress modality (22). In principle, the gene regulatory apparatus can integrate a variety of stress related signals over time producing a more stable and robust signal for measurement.

The stress-sensitive gene expression index described here appears to be as sensitive as most behavioral tests currently used to measure chronic stress exposure. Isolation housing is established as a stressor in rats (29-31) but its effects are not always apparent using standard behavioral assays (21, 32) and results from different studies can be contradictory (31). The SSGE index could clearly distinguish the social isolation group (rats housed singly) from the control group (rats housed three per cage) (Figure 2C), suggesting that the assay is at least as sensitive as typical behavioral tests.

Many of the genes found to be differentially expressed in the adrenal gland following stress exposure in this study have previously been reported to be stress-sensitive (33, 34). All of the genes included in the SSGE index (Table 1) have been implicated in the response to stress in one or more tissues (35-43). Currently, however, relatively little is known about the specific role that these genes might play in mediating the response to stress in the adrenal gland. Phenylalanine Hydroxylase (Pah) catalyzes the irreversible conversion of L-phenylalanine into L-tyrosine, an essential step in the catecholamine biosynthesis pathway (44). However, it is generally believed that tyrosine synthesis occurs predominantly in the liver, with only the downstream catecholamine synthesis steps occurring in the adrenal medulla (10). Three genes (Thrsp, Scd and Cd36) are involved in fatty acid metabolism. The adrenal gland has a high content of lipids, whose main function is to serve as precursors for steroid hormone biosynthesis in the cortex (45). Changes in expression of genes implicated in lipid metabolism and transport in the adrenal gland may indicate a long-term effect of stress on steroid hormone biosynthesis. Expression of Cadherin8 (Cdh8) is decreased in the prefrontal cortex in response to stress and it has been hypothesized that this may alter tissue plasticity to favor adaptive synapse remodeling (43). The Na/H exchanger (Slc9a3) uses an inward sodium ion gradient to expel acids from the cell (46). Its role in sodium and pH homeostasis has been studied mainly in the kidney and intestine and its function in adrenal gland tissue has not been characterized.

**Table 1.** SSGE Index Genes and Their Potential Role in Stress.

| **Protein Name** | **Gene Symbol** | **Specific Function** | **Potential Role in Stress Response** |
| --- | --- | --- | --- |
| Phenylalanine Hydroxylase | Pah | Converts Phenylalanine into Tyrosine | Catecholamine biosynthesis |
| Sodium-Hydrogen Antiporter 3 | Slc9a3 | Imports one Na<sup>+</sup> in the cell/exports one H<sup>+</sup> | unknown |
| Thyroid Hormone Responsive gene | Thrsp | Regulator of lipid metabolism | Altered fatty acid metabolism in adrenal gland |
| Stearoyl-CoA desaturase | Scd | Biosynthesis of monounsaturated fatty acids | Altered fatty acid metabolism in adrenal gland |
| Cadherin 8 | Cdh8 | Mediates cell-cell adhesion | Adaptive changes in tissue plasticity |
| Cluster of Differentiation 36 | Cd36 | Fatty Acid Transport | Altered fatty acid metabolism in adrenal gland |

Molecular diagnostic measures of traumatic stress exposure in humans are an important and realistic goal (2). Robust tests would be of considerable value in predicting the risk of developing stress related disorders such as PTSD and stress-induced depression. This report suggests that one viable approach towards the development of such a test is an index based on those changes in gene regulatory function that are universal across a range of stress modalities. The stress-sensitive gene regulatory index performs as required for such a diagnostic test, since it responds in a dose-dependent way to stress exposure, is independent of the modality of stress exposure, and is precise, even while relying on a relatively small number of measured variables. A practical diagnostic test in humans would necessarily use tissues that can be easily biopsied such as peripheral blood mononuclear or buccal cells, which also show stress-induced changes in gene regulatory function (5, 6, 47, 48).

## Acknowledgements

The project was supported by a VA Merit Review Grant, a Carol M. Baldwin Award and the Institute of Molecular Cardiology. We would like to thank Yezdan Pece, Yuyan Huang, Katelyn Neuman and Arun Nallainathan for their assistance with the experiments.

## References

1. Yehuda R, Hoge CW, McFarlane AC, Vermetten E, Lanius RA, Nievergelt CM, et al. (2015): Post-traumatic stress disorder. Nat Rev Dis Primers. 1:15057.

2. Lehrner A, Yehuda R (2014): Biomarkers of PTSD: military applications and considerations. Eur J Psychotraumatol. 5.

3. Bagot RC, Cates HM, Purushothaman I, Lorsch ZS, Walker DM, Wang J, et al. (2016): Circuit-wide Transcriptional Profiling Reveals Brain Region-Specific Gene Networks Regulating Depression Susceptibility. Neuron. 90:969–983.

4. Muhie S, Gautam A, Meyerhoff J, Chakraborty N, Hammamieh R, Jett M (2015): Brain transcriptome profiles in mouse model simulating features of post-traumatic stress disorder. Mol Brain. 8:14.

5. Bierhaus A, Wolf J, Andrassy M, Rohleder N, Humpert PM, Petrov D, et al. (2003): A mechanism converting psychosocial stress into mononuclear cell activation. Proc Natl Acad Sci U S A. 100:1920–1925.

6. Muhie S, Gautam A, Chakraborty N, Hoke A, Meyerhoff J, Hammamieh R, et al. (2017): Molecular indicators of stress-induced neuroinflammation in a mouse model simulating features of post traumatic stress disorder. Transl Psychiatry. 7:e1135.

7. Davidson EH, Rast JP, Oliveri P, Ransick A, Calestani C, Yuh CH, et al. (2002): A genomic regulatory network for development. Science. 295:1669–1678.

8. Bali A, Jaggi AS (2015): Preclinical experimental stress studies: protocols, assessment and comparison. Eur J Pharmacol. 746:282–292.

9. Patchev VK, Patchev AV (2006): Experimental models of stress. Dialogues Clin Neurosci. 8:417–432.

10. Kvetnansky R, McCarty R (2010): Adrenal Medulla In: Fink G, editor. Stress Science: Neuroendocrinology. San Diego: Elsevier, pp 261–268.

11. Vinson GP, Whitehouse BJ, Hinson JP (2010): Adrenal Cortex In: Fink G, editor. Stress Science: Neuroendocrinology. San Diego: Elsevier, pp 137–145.

12. Liao Y, Smyth GK, Shi W (2013): The Subread aligner: fast, accurate and scalable read mapping by seed and vote. Nucleic Acids Res. 41:e108.

13. Liao Y, Smyth GK, Shi W (2014): featureCounts: an efficient general purpose program for assigning sequence reads to genomic features. Bioinformatics. 30:923–930.

14. Ruijter JM, Pfaffl MW, Zhao S, Spiess AN, Boggy G, Blom J, et al. (2013): Evaluation of qPCR curve analysis methods for reliable biomarker discovery: bias, resolution, precision, and implications. Methods. 59:32–46.

15. Walf AA, Frye CA (2007): The use of the elevated plus maze as an assay of anxiety related behavior in rodents. Nat Protoc. 2:322–328.

16. Pellow S, Chopin P, File SE, Briley M (1985): Validation of open:closed arm entries in an elevated plus-maze as a measure of anxiety in the rat. J Neurosci Methods. 14:149–167.

17. Davis M (1989): Sensitization of the acoustic startle reflex by footshock. Behav Neurosci. 103:495–503.

18. Valsamis B, Schmid S (2011): Habituation and prepulse inhibition of acoustic startle in rodents. J Vis Exp. e3446.

19. Benjamini Y, Hochberg Y (1995): Controlling the False Discovery Rate: A Practical and Powerful Approach to Multiple Testing. Journal of the Royal Statistical Society Series B (Methodological). 57:289–300.

20. R_Core_Team (2017): R: A language and environment for statistical computing. Vienna, Austria.: R Foundation for Statistical Computing.

21. Nakayasu T, Ishii K (2008): Effects of pair-housing after social defeat experience on elevated plus-maze behavior in rats. Behav Processes. 78:477–480.

22. Dickens MJ, Romero LM (2013): A consensus endocrine profile for chronically stressed wild animals does not exist. Gen Comp Endocrinol. 191:177–189.

23. Wilson MA, Grillo CA, Fadel JR, Reagan LP (2015): Stress as a one-armed bandit: Differential effects of stress paradigms on the morphology, neurochemistry and behavior in the rodent amygdala. Neurobiol Stress. 1:195–208.

24. Wilker S, Pfeiffer A, Kolassa S, Koslowski D, Elbert T, Kolassa IT (2015): How to quantify exposure to traumatic stress? Reliability and predictive validity of measures for cumulative trauma exposure in a post-conflict population. Eur J Psychotraumatol. 6:28306.

25. Neuner F, Schauer M, Karunakara U, Klaschik C, Robert C, Elbert T (2004): Psychological trauma and evidence for enhanced vulnerability for posttraumatic stress disorder through previous trauma among West Nile refugees. BMC Psychiatry. 4:34.

26. Mollica RF, McInnes K, Poole C, Tor S (1998): Dose-effect relationships of trauma to symptoms of depression and post-traumatic stress disorder among Cambodian survivors of mass violence. Br J Psychiatry. 173:482–488.

27. Kolassa I-TE, Verena; Eckart, Cindy; Kolassa, Stephan; Onyut, Lamaro P.; Elbert, Thomas (2010): Spontaneous remission from PTSD depends on the number of traumatic event types experienced. Psychological Trauma: Theory, Research, Practice, and Policy. 2:169–174.

28. Marti O, Armario A (1998): Anterior pituitary response to stress: time-related changes and adaptation. Int J Dev Neurosci. 16:241–260.

29. Weintraub A, Singaravelu J, Bhatnagar S (2010): Enduring and sex-specific effects of adolescent social isolation in rats on adult stress reactivity. Brain Res. 1343:83–92.

30. Dronjak S, Gavrilovic L, Filipovic D, Radojcic MB (2004): Immobilization and cold stress affect sympatho-adrenomedullary system and pituitary adrenocortical axis of rats exposed to long-term isolation and crowding. Physiol Behav. 81:409–415.

31. Burke AR, McCormick CM, Pellis SM, Lukkes JL (2017): Impact of adolescent social experiences on behavior and neural circuits implicated in mental illnesses. Neurosci Biobehav Rev.

32. Bourke CH, Neigh GN (2011): Behavioral effects of chronic adolescent stress are sustained and sexually dimorphic. Horm Behav. 60:112–120.

33. Fedoseeva LA, Klimov LO, Ershov NI, Alexandrovich YV, Efimov VM, Markel AL, et al. (2016): Molecular determinants of the adrenal gland functioning related to stress-sensitive hypertension in ISIAH rats. BMC Genomics. 17:989.

34. Murani E, Ponsuksili S, D’Eath RB, Turner SP, Evans G, Tholking L, et al. (2011): Differential mRNA expression of genes in the porcine adrenal gland associated with psychosocial stress. J Mol Endocrinol. 46:165–174.

35. Ryazanova MA, Fedoseeva LA, Ershov NI, Efimov VM, Markel AL, Redina OE (2016): The gene-expression profile of renal medulla in ISIAH rats with inherited stress-induced arterial hypertension. BMC Genet. 17:151.

36. Imumorin IG, Dong Y, Zhu H, Poole JC, Harshfield GA, Treiber FA, et al. (2005): A gene environment interaction model of stress-induced hypertension. Cardiovasc Toxicol. 5:109–132.

37. Liu X, Strable MS, Ntambi JM (2011): Stearoyl CoA desaturase 1: role in cellular inflammation and stress. Adv Nutr. 2:15–22.

38. Namboodiri MA, Ramasarma T (1975): Effect of environmental stress of low pressure on tyrosine aminotransferase and phenylalanine 4 hydroxylase activities in the rat. Biochem J. 150:263–268.

39. Putri M, Syamsunarno MR, Iso T, Yamaguchi A, Hanaoka H, Sunaga H, et al. (2015): CD36 is indispensable for thermogenesis under conditions of fasting and cold stress. Biochem Biophys Res Commun. 457:520–525.

40. Okamura DM, Pennathur S, Pasichnyk K, Lopez Guisa JM, Collins S, Febbraio M, et al. (2009): CD36 regulates oxidative stress and inflammation in hypercholesterolemic CKD. J Am Soc Nephrol. 20:495–505.

41. Chuang JC, Cui H, Mason BL, Mahgoub M, Bookout AL, Yu HG, et al. (2010): Chronic social defeat stress disrupts regulation of lipid synthesis. J Lipid Res. 51:1344–1353.

42. Lebsack TW, Fa V, Woods CC, Gruener R, Manziello AM, Pecaut MJ, et al. (2010): Microarray analysis of spaceflown murine thymus tissue reveals changes in gene expression regulating stress and glucocorticoid receptors. J Cell Biochem. 110:372–381.

43. de Azeredo LA, Wearick Silva LE, Viola TW, Tractenberg SG, Centeno Silva A, Orso R, et al. (2017): Maternal separation induces hippocampal changes in cadherin 1 (CDH1) mRNA and recognition memory impairment in adolescent mice. Neurobiol Learn Mem. 141:157–167.

44. Flydal MI, Martinez A (2013): Phenylalanine hydroxylase: function, structure, and regulation. IUBMB Life. 65:341–349.

45. Walther TC, Farese RV, Jr. (2012): Lipid droplets and cellular lipid metabolism. Annu Rev Biochem. 81:687–714.

46. Donowitz M, Li X (2007): Regulatory binding partners and complexes of NHE3. Physiol Rev. 87:825–872.

47. Mathews HL, Konley T, Kosik KL, Krukowski K, Eddy J, Albuquerque K, et al. (2011): Epigenetic patterns associated with the immune dysregulation that accompanies psychosocial distress. Brain Behav Immun. 25:830–839.

48. Non AL, Hollister BM, Humphreys KL, Childebayeva A, Esteves K, Zeanah CH, et al. (2016): DNA methylation at stress-related genes is associated with exposure to early life institutionalization. Am J Phys Anthropol. 161:84–93.

